# A combination of *EWSR1-FLI1* with loss of one *EWSR1* allele promotes damage in the mitotic spindle and sensitization to microtubule-destabilizing agent

**DOI:** 10.64898/2026.06.15.732411

**Authors:** Evan T Schulz, Mizuki Azuma

**Affiliations:** University of Kansas, Molecular Biosciences, Lawrence KS, Unites States

**Author notes:** **Correspondence:** Corresponding Author: Mizuki Azuma.

## Abstract

Microtubule-destabilizing agents (MDAs) and microtubule-stabilizing agents (MSAs) are commonly used chemotherapeutic agents due to its activity to induce cell death by compromising the dynamics of spindles during mitosis. Ewing sarcoma, the second most common pediatric bone cancer, is known to selectively respond to MDAs as a first-line treatment, but not to MSAs. Ewing sarcoma cells carry an aberrant *EWSR1-FLI1* fusion gene and only one wildtype *EWSR1* allele. To investigate the origin of this MDA sensitivity, we used an (*AID-EWSR1/wt: EWSR1-FLI1-mCherry/wt*) cell line that enables conditional induction of EWSR1-FLI1 expression (Tet-On system) and EWSR1 knockdown derived from one *EWSR1* allele (auxin-degron system). A combination of EWSR1-FLI1 expression and EWSR1 knockdown induces apoptosis upon nocodazole treatment, a type of MDA. Our study revealed that the mitotic spindles of Ewing sarcoma cells (A673, RD-ES and SK-ES1) contain elevated levels of tubulin damage, visualized with GTP-tubulin, compared to mesenchymal stem cells (MSC). Consistently, the combination of EWSR1-FLI1 expression and EWSR1 knockdown in (*AID-EWSR1/wt: EWSR1-FLI1-mCherry/wt*) cell line also leads to an increased incidence of damage in mitotic spindles. Together, we propose that the sensitivity of Ewing sarcoma cells is derived from the increased levels of damage to mitotic spindles caused by EWSR1-FLI1 expression and EWSR1 knockdown.

## INTRODUCTION

Vinka alkaloid derived MDAs, including Vincristine, are a common class of chemotherapy drugs that have been used for cancer treatment since the 1960’s [1]. The clinical effectiveness of MDAs derives from their tubulin binding activity that causes mitotic spindle destabilization and dynamism defects, compromising mitotic progression and inducing cell death [2, 3]. MDAs (e.g. Vincristine) are essential to the chemotherapeutic treatment of Ewing sarcoma, a pediatric bone cancer, validated by decades of demonstrated sensitivity and clinical effectiveness [2, 3]. In contrast, microtubule-stabilizing agents (MSAs, e.g. paclitaxel) are less effective for Ewing sarcoma patients as a first-line treatment [4, 5]. While both drug classes impair mitosis, it is unclear why Ewing sarcoma is more sensitive to MDAs than MSAs. Therefore, our study aims to fill this knowledge gap by elucidating the etiology of Ewing sarcoma’s MDA sensitivity.

All Ewing sarcoma patients express an aberrant fusion gene that contains the N-terminus coding sequence of EWSR1 and the C-terminus coding sequence of one of the *ETS* genes (*FLI1, ERG, ETV1* and *ETV5*) [6-8]. Among these, *EWSR1-FLI1* is the predominant fusion gene observed in Ewing sarcoma patients at approximately 85% [6, 9]. Our previous studies demonstrated a unique function of EWSR1-FLI1, in which it induces mitotic dysfunction and aneuploidy in human cells and zebrafish [10-13]. We also discovered that EWSR1-FLI1 interacts with EWSR1 and induces mitotic dysfunction [14]. One possible cause of mitotic dysfunction in Ewing sarcoma cells is damage to mitotic spindles, an important type of microtubule defect. This damage results from the emergence of gaps within the spindle lattice due to the detachment of α-tubulin and β-tubulin dimers. Microtubule damage may enhance the sensitivity spindle fibers to MDAs by compromising fiber stability, dynamism, increasing availability of binding sites for various MDAs or inducing lattice deacetylation [15-20]. Damage is repaired and stabilized by the incorporation of GTP-tubulin, resulting in islands of GTP-tubulin within the microtubule lattice [21]. While complete repair of spindle damage can ameliorate tubulin lattice stability, GTP-tubulin is often rapidly hydrolyzed to GDP-tubulin to maintain proper lattice conformation, fiber dynamism, motor protein translocation, and microtubule-associated protein (MAP) interactions [21-27]. These results suggest that high levels of GTP-tubulin incorporation in microtubules indicate areas of compromised dynamism in mitotic spindles. While EWSR1 interacts with α-tubulin, and is reported to stabilize the non-mitotic tubulin structure, it is unknown how EWSR1 and EWSR1-FLI1 regulate the mitotic spindle structure [28]. Here, we propose that the sensitization of cell expressing EWSR1-FLI1 and knockdown of EWSR1 results from an increase in tubulin damage that enhances the MDA mechanism of action.

## MATERIAL AND METHODS

### Cell culture and induction of EWSR1 knockdown and EWSR1-FLI1 expression

The Ewing sarcoma cells (A673, RD-ES and SK-ES1) and MSC were cultured following the protocol provided by the American Type Culture Collection (ATCC). The (*AID-EWSR1/wt;EWSR1FLI1-mCherry/wt*) DLD-1 cell line (OSTIR1-7-21-38) was maintained in McCoy’s media supplemented with 10% Tet-free Fetal Bovine Serum (FBS), 1 μg/mL Blasticidin (Invivogen, #ant-bl), 200 μg/mL Hygromycin B Gold (Invivogen, #ant-hg), and 1ug/ml puromycine [12, 13]. To induce EWSR1 knockdown resulting from one *EWSR*1 allele and/or EWSR1-FLI1 expression, the cells were subjected to four conditions over either 24hrs or 7 days: control (treated with DMSO), EWSR1 knockdown (treated with 1mM Auxin (AUX)), EWSR1-FLI1 expression (treated with 1ug/ml of doxycycline (DOX)), and EWSR1 knockdown/EWSR1-FLI1 expression (treated with 1mM AUX and 1ug/ml of DOX).

### Apoptosis Assay

The apoptotic cells were visualized with Annexin V Alexa Flour 568 by following the protocol supplied by the company (Invitrogen, Cat: A13202). The cells were photo-documented with EVOS imaging systems (Thermo Fisher Scientific), and the percentage of apoptotic cells was calculated.

### Immunocytochemistry and Tubulin Damage assay

The cells were permeabilized, stained with hMB11 and anti-human IgG secondary antibodies, fixed, and co-labeled, according to the protocol previously established by with minor modification [29]. For all staining, washing, and α-tubulin co-labeling steps, PEMG (80mM PIPES, 0.5mM EDTA, 2mM MgCl2, 10% Glycerol) was used. To visualize mitotic spindles, EWSR1 and EWSR1-FLI1-mCherry, the cells were treated with rabbit anti-α Tubulin (Sigma-Aldrich, Cat: 11224-1-AP, 1:500 dilution), mouse anti-EWSR1 (Abnova, Cat: H00002130-M01, 1:500 dilution) and chicken anti-mCherry (LSBio, Cat: LS-C204825, 1:500 dilution) for 3 hours at room temperature. The cells were subjected to secondary antibodies Goat anti-Rabbit IgG Alexa Fluor 405nm (Invitrogen, Cat: A-31556 1:250 dilution), Goat anti-Mouse IgG Alexa Fluor 488 (Invitrogen, Cat: A-32723, 1:250 dilution), and Goat anti-Chicken IgG Alex Fluor 568 (Invitrogen, Cat: A-11041 1:250 dilution) at room temperature for 3 hours. The cells were washed three times with 4°C PEM-G for 5 minutes at room temperature and mounted onto slides with antifade mounting medium (VECTASHIELD, Cat: H-1000).

To visualize damage in the mitotic spindle lattice, cells were visualized with anti-Tubulin-GTP (Adipogen, Cat: AG-27B-0009-C100, 1:5000 dilution) and anti-Human AlexFlour 488nm (Invitrogen, Cat: A-11013, 1:1,000 dilution) following the previous report, and mounted onto slides with antifade mounting medium (VECTASHIELD, Cat: H-1000) [29].

### Documentation of cell images

Cell images were documented by capturing Z-sections at 0.2 µm intervals using spinning disk confocal microscopy (Nikon, Crest Optics X-Light V3 spinning disk confocal on Eclipse Ti2-E), followed by analysis with an image software FIJI.

### Statistics

Standard deviation (SD) is shown as error bars for each graph. Statistical confidence was defined at *p* < 0.05 by ANOVA two-way analysis, followed by the post-hoc test using Tukey’s Honesty Significant Difference (HSD).

## RESULTS

While sensitivity to MDAs has been established for Ewing sarcoma cells, understanding of the cause of this tumor type-specific sensitivity has been limited. To confirm the effect of MDAs, we employed nocodazole, which depolymerizes spindles by interacting with monomer β-tubulin [2, 3]. Ewing sarcoma cells (A673, RD-ES and SK-ES1) and MSC were treated with nocodazole for 24 hours, and the incidence of apoptosis was measured by Annexin V staining. The apoptosis rate of the nocodazole-treated Ewing sarcoma cells (A673, RD-ES and SK-ES1) was significantly higher than that of MSC, supporting the previous observation of Ewing sarcoma sensitivity to MDAs (**Fig 1A**) [5]. Importantly, nocodazole treatment in MSC did not induce a significant increase in apoptosis, suggesting a nocodazole sensitivity specific to Ewing sarcoma (**Fig 1A**).

**Fig 1.**
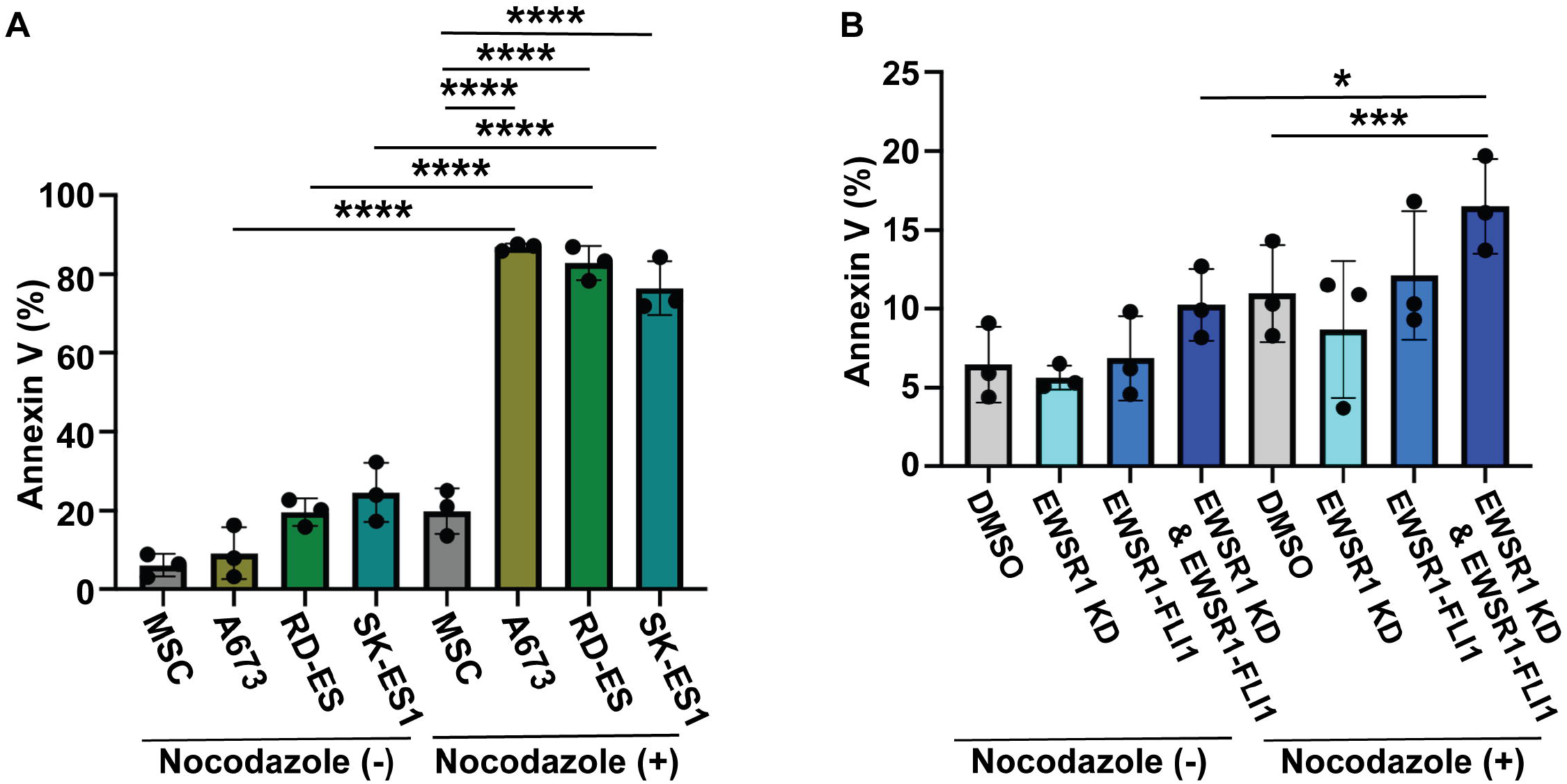
Nocodazole treatment induces an increased incidence of apoptosis in Ewing sarcoma cells and DLD-1 cells with EWSR1 knockdown and EWSR1-FLI1 expression. **A**Percentages of cells undergoing apoptosis were visualized by Annexin V staining in Ewing sarcoma cells (A673, RD-ES, and SK-ES1) and Mesenchymal Stem Cells (MSC). **B**. Percentages of apoptotic cells were visualized by Annexin V staining in four samples induced in (*AID-EWSR1/wt: EWSR1-FLI1-mCherry/wt*) DLD-1 cells: control (DMSO), EWSR1 knockdown (AUX), EWSR1-FLI1 (DOX) and EWSR1 knockdown/EWSR1-FLI1 (AUX/DOX). A minimum of fifty cells were scored for each experiment, and the experiments were repeated three times for all experiments (**A** and **B**). Nocodazole (-): cells treated without Nocodazole, Nocodazole (+): cells treated with 332nM Nocodazole. *: P<0.05, ***: P<0.001, ****: P<0.0001

To assess the effect of *EWSR1-FLI1* and the loss of one *EWSR1* allele, we utilized a (*AID-EWSR1/wt: EWSR1-FLI1-mCherry/wt*) cell line that was originally generated using a CRISPR/Cas9 system [30]. This cell line enables the conditional expression of EWSR1-FLI1 using the Tet-on system, and conditional degradation of EWSR1 derived from one *EWSR1* allele using the auxin-degron system. This conditional cell line was generated in the DLD-1 cell line, as it does not express EWSR1-FLI1 and has a near-diploid karyotype [31, 32]. The cells were treated with AUX and/or DOX to degrade EWSR1 derived from one *EWSR1* allele and induce EWSR1-FLI1 expression respectively. Concurrently, these treatment groups were treated with or without nocodazole for twenty-four hours. These eight samples were subjected to Annexin V staining, and their rates of apoptosis were calculated. As a result, the cells that underwent treatment to induce EWSR1-FLI1 expression and EWSR1 knockdown displayed significantly higher apoptosis compared to the control (**Fig 1B**). These results suggest that the combination of EWSR1-FLI1 expression and EWSR1 knockdown is required for the acquisition of the sensitivity to nocodazole.

Since EWSR1-FLI1 and EWSR1 share the same N-terminus of EWSR1, there is a technical challenge in distinguishing between the EWSR1-FLI1 and EWSR1 using immunocytochemistry. To investigate whether EWSR1-FLI1 and EWSR1 localize at the mitotic spindle, we utilized the (*AID-EWSR1/wt: EWSR1-FLI1-mCherry/wt*) cells. The cell was subjected to immunocytochemistry using an anti-EWSR1 antibody to visualize AID-EWSR1 (green), anti-mCherry to visualize EWSR1-FLI1-mCherry (red) and anti-α tubulin antibody to visualize mitotic spindle (blue). These cells were documented using a confocal microscope. Single Z-section images of the cells were analyzed with the image software Fiji to study the colocalization of combinations of EWSR1 and α-tubulin (Person’s coefficient: 0.76+/-0.04), EWSR1-FLI1 and α-tubulin (Person’s coefficient: 0.80+/-0.04), and EWSR1-FLI1 and EWSR1 (Person’s coefficient: 0.77+/-0.06) (**Fig S1A, B and C**). The result suggests that EWSR1-FLI1 and EWSR1 colocalize on mitotic spindles in the (*AID-EWSR1/wt: EWSR1-FLI1-mCherry/wt*) cells.

Previous studies have shown that EWSR1 interacts with α-tubulin, and stabilizes microtubules during the non-mitotic phase [28]. Based on this, we hypothesized that the combination of EWSR1-FLI1 expression and EWSR1 knockdown would induce damage in mitotic spindles. The (*AID-EWSR1/wt: EWSR1-FLI1-mCherry/wt*) cells were treated with AUX and/or DOX for 24 hours (hrs) and 7 days, and then subjected to immunocytochemistry using anti-GTP-Tubulin (green) and anti-α-tubulin (blue) antibodies (**Fig 2A and B**). The intensity of the signals for anti-α-tubulin and GTP-tubulin was measured, and the signals of GTP-tubulin were normalized against the signals of α-tubulin (**Fig 2C and D**). The signals of GTP-tubulin were significantly higher in the both cells with EWSR1 knockdown (AUX) and EWSR1-FLI1 expression (DOX) compared to the control (DMSO) for 24 hrs and 7 days (**Fig 2C and D**). Moreover, the combination of EWSR1 knockdown and EWSR1-FLI1 expression (AUX/DOX) was higher than either EWSR1 knockdown (AUX) or EWSR1-FLI1 expression (DOX) alone, when compared to the control (DMSO) after 24 hrs and 7 days (**Fig 2C and D**). The result suggests that combination of EWSR1 knockdown and EWSR1-FLI1 expression induces damage in mitotic spindles.

**Fig 2.**
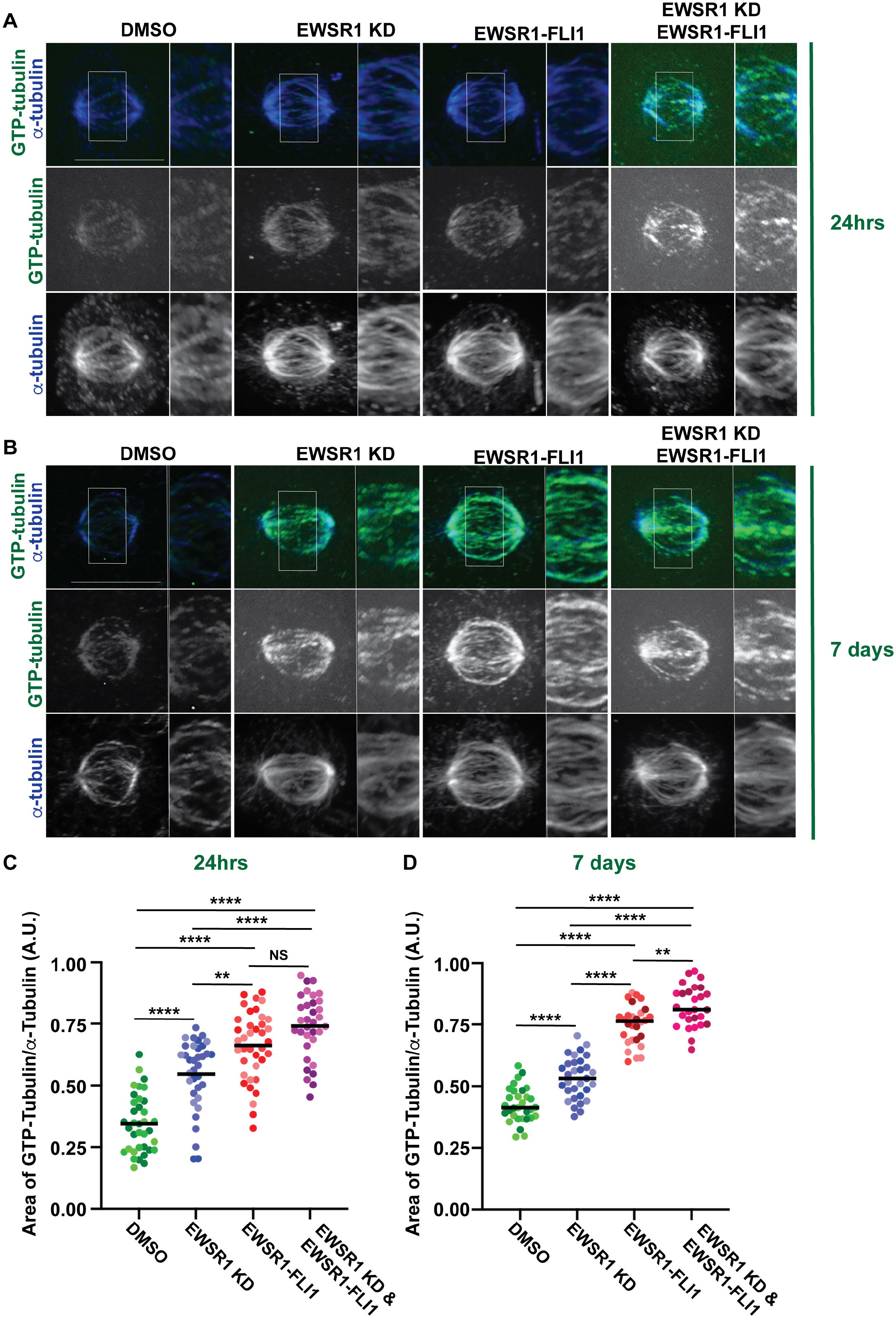
A combination of EWSR1 knockdown and EWSR1-FLI1 expression results in the induction of damage to microtubules. **A and B** Representative images of metaphase spindles of (*AID-EWSR1/wt: EWSR1-FLI1-mCherry/wt*) DLD-1 cells. The cells were visualized with anti-α tubulin (blue) and anti-GTP tubulin (green) obtained from control (DMSO), EWSR1 knockdown (AUX), EWSR1-FLI1 expression (DOX), and EWSR1 knockdown/EWSR1-FLI1 expression (AUX/DOX). Scale bar = 10µm. **C and D**. The area of GTP-tubulin signals was normalized against the area of α-tubulin. A minimum twenty cells per sample were quantified, and the experiment was repeated for n=3 experiments. (**A and C**. treated with AUX and/or DOX for 24hrs, and **B and D**. treated with AUX and/or DOX for 7 days). **: P<0.01, ****: P<0.0001, NS: Non-Significant.

To investigate whether Ewing sarcoma cells exhibit the same damage in mitotic spindles, we performed immunocytochemistry using anti-GTP-Tubulin (green) and anti-α-tubulin (blue) antibodies in Ewing sarcoma cells (A673, RD-ES and SK-ES1). We then compared its intensity of staining with that of MSC, which is one of the proposed cell of origin of Ewing sarcoma (**Fig 3A**) [33]. The intensity of GTP-tubulin in Ewing sarcoma cells was significantly higher than in MSC (**Fig 3B**). Note that the intensity of GTP-tubulin was significantly higher in RD-ES and SK-ES1 compared to A673 with an unknown mechanism. These results indicate the level of spindle damage is higher in Ewing sarcoma than in MSCs.

**Fig 3.**
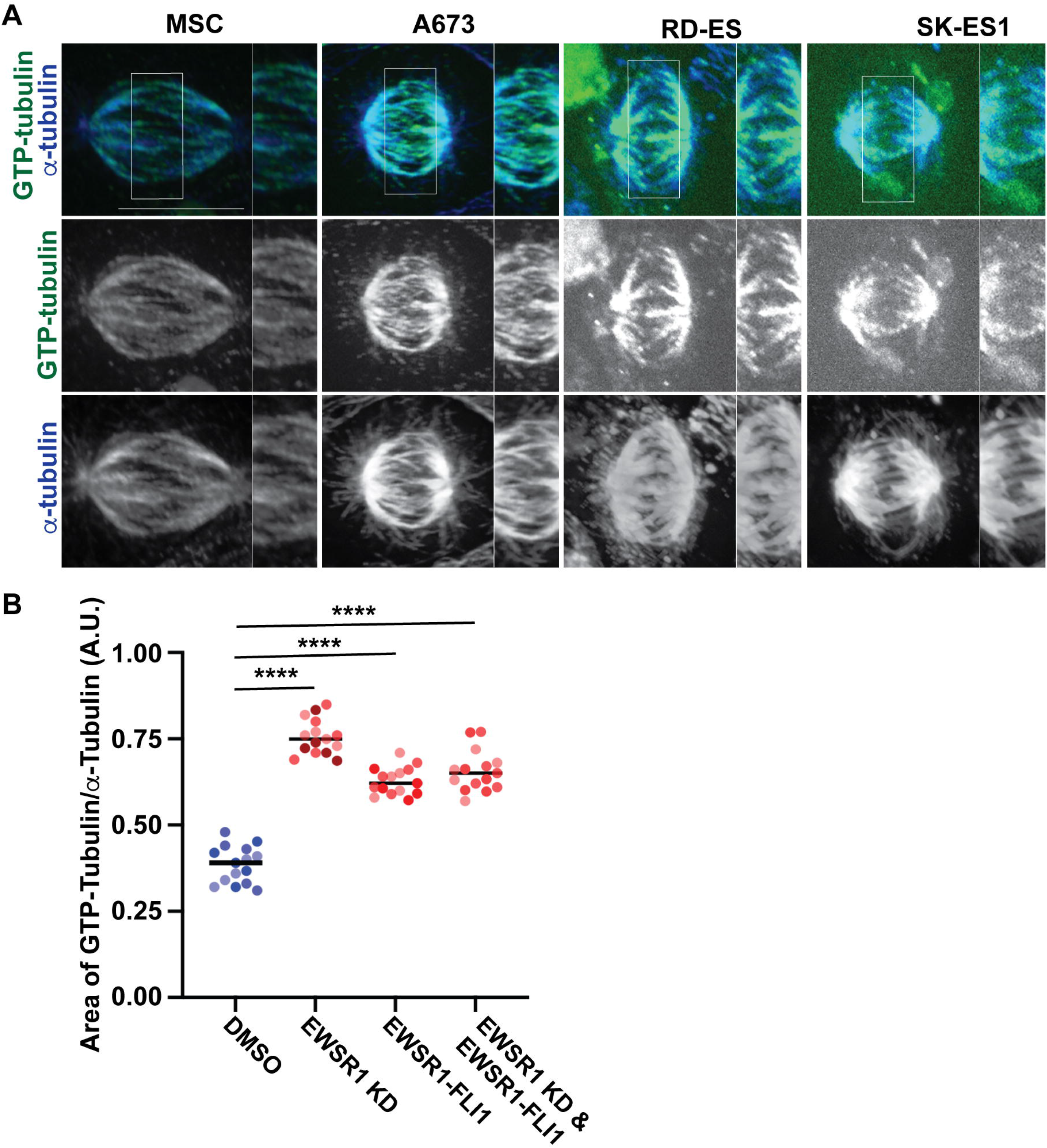
Ewing sarcoma cells have higher levels of damage to microtubules than MSC. Representative images of metaphase spindles visualized with anti-α tubulin (blue) and anti-GTP tubulin (green) in MSC, A673, RD-ES, and SK-ES1 cells. The intensity of the signals from a minimum of five cells was quantified from n=3 experiments. ****: P<0.0001). Scale bar = 10µm.

## DISCUSSION

This study aimed to investigate the tumor type-specific cause of MDA sensitivity observed in Ewing sarcoma patients. We discovered that Ewing sarcoma cells exhibit elevated levels of GTP-tubulin in their mitotic spindles, suggesting an increased level of damage to the mitotic spindles compared to MSC. Ewings sarcoma cells also undergo a higher rate of apoptosis under nocodazole treatment compared to MSCs. Furthermore, mimicking the genotype of Ewing sarcoma by knocking down EWSR1 and expressing EWSR1-FLI1 conditionally in DLD-1 cells increased levels of GTP-tubulin in mitotic spindles and the rate of apoptosis under nocodazole treatment. These data suggest that the combination of EWSR1 knockdown and EWSR1-FLI1 expression is sufficient to simultaneously induce mitotic spindle damage and sensitize cells to nocodazole microtubule disruption. Together, we demonstrate that mitotic spindle damage caused by Ewing sarcoma-specific aberrations may contribute to the tumor type’s sensitivity to MDAs, like nocodazole.

Damage to the tubulin lattice induces mechanical stress and conformational deformations across the microtubule lattice, causing loss of fiber stability, stiffness, and dynamism, as well as, altering MAP interactions and motor protein trafficking [15, 17, 23, 34-36]. To further understand how the stability of mitotic spindles is compromised in Ewing sarcoma, mitotic spindle formation rate, dynamism, and post translational modifications (PTM) need to be characterized by the following means in Ewing sarcoma cells. First, to study the rate of spindle formation in Ewing sarcoma cells, the rate of microtubule polymerization should be quantified. Second, dynamic instability should be quantified by measuring the rate of microtubule switching between polymerization and depolymerization. The damaged tubulin lattice is unstable and remains unstable until the gaps in the lattice are replaced with GTP-tubulin. Therefore, it is plausible that majority of mitotic spindles in Ewing sarcoma cells are destabilized. If the rate of microtubule formation or dynamic instability are reduced in Ewing sarcoma, it will provide evidence of spindle destabilization and support our observation that Ewing sarcoma is sensitized to MDAs. Additionally, PTM status of mitotic spindles can enhance their stability. A previous study demonstrated that the EWSR1 promotes acetylation of spindles through modulation of Histone Deacetylase (HDAC) 6 [28]. Therefore, spindle acetylation should be quantified in Ewing sarcoma cells as it is plausible that the loss of the *EWSR1* allele affects the rate of spindle acetylation and destabilizes mitotic spindles.

In this study, we observed that the levels of GTP-tubulin in RD-ES and SK-ES1 cells were higher than in A673 cells, suggesting a variation of the degree in the tubulin damage among Ewing sarcoma patients. One possible explanation for the differences among patient cells is the presence of polymorphisms in α-tubulin and β-tubulin genes. For example, The T238A mutation in human β-tubulin stabilizes tubulin by altering the rate of GTP-uncoupling through its conformational change within the microtubule [37]. The H283Y mutation in α-tubulin was proposed to stabilize tubulin filaments by promoting the interaction between longitudinal interdimer, intradimer and lateral interactions between protofilaments [38]. Therefore, polymorphisms in tubulin genes could affect MDA sensitivity or resistance by directly modifying spindle stability. Likewise, it will be important to subclassify the types of polymorphisms that are not sensitive to microtubule depolymerizing agents. Identification and classification of tubulin gene polymorphisms in patients could predict the drug efficacy, indicate resistance formation, and inform treatment strategies.

## ACKNOWLEDGEMENT

The study was supported by the Braden’s Hope Foundation Pilot Grant, Massman’s family research fund, Alan B Slifka Foundation Research Fund, and University of Kansas (KU) General Research Fund (GRF).

## FIGURE LEGENDS

**Fig S1.**
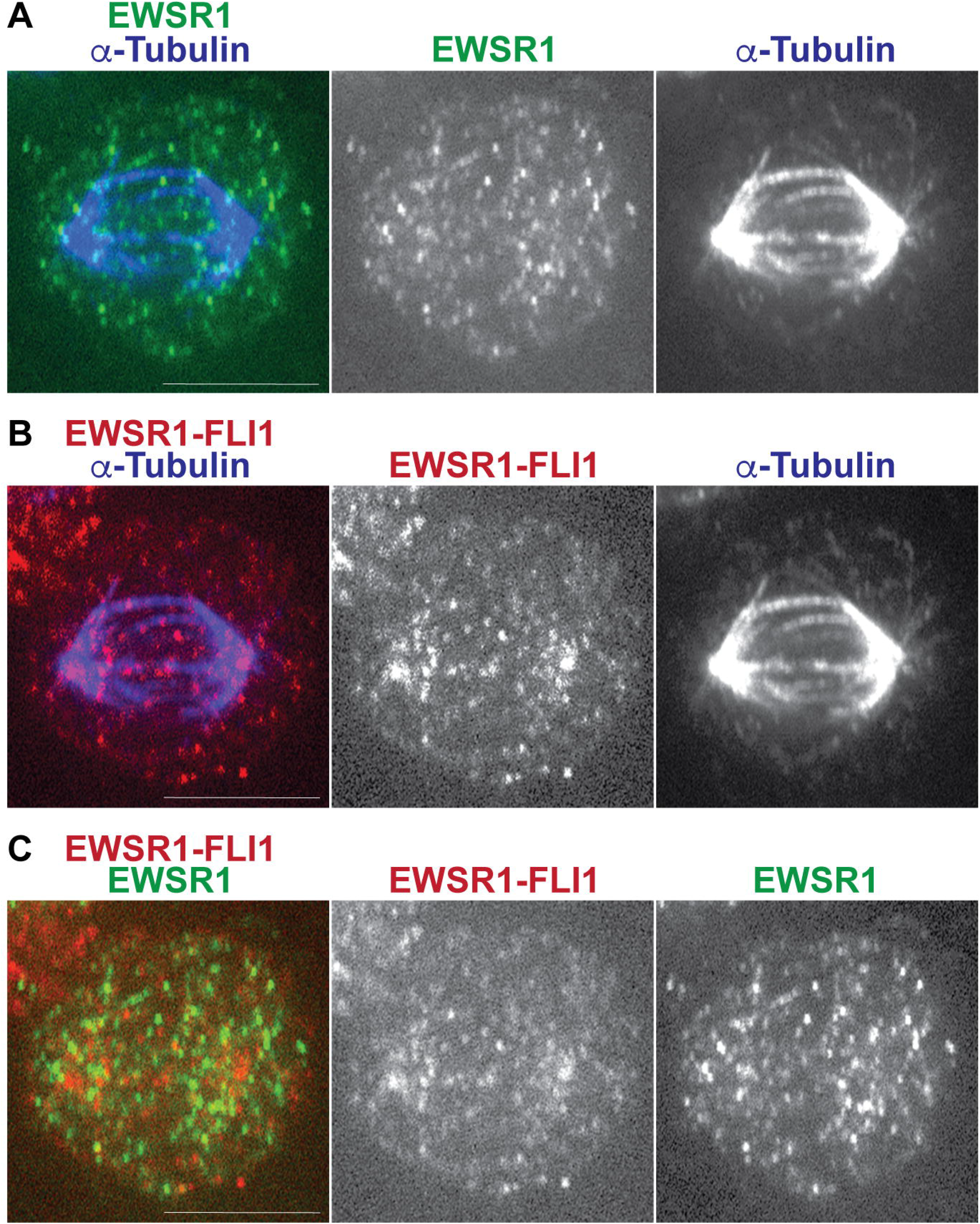
Co-localization of EWSR1 and EWSR1-FLI1 during mitosis. Single Z-section plane immunofluorescence image of a metaphase (*AID-EWSR1/wt: EWSR1-FLI1-mCherry/wt*) DLD-1 cell expressing ectopic EWSR1-FLI1 (DOX). α tubulin, EWSR1, and EWSR1-FLI1 were visualized using anti-α tubulin, anti-EWSR1, and anti-mCherry antibodies, respectively (**A**. EWSR1 and α tubulin, **B**. EWSR1-FLI1 and α tubulin, and **C**. EWSR1 and EWSR1-FLI1). Scale bar = 10 µm.

## REFERENCES

1 Cutts, J. H., Beer, C. T., Noble, R. L. 1960 Biological properties of Vincaleukoblastine, an alkaloid in Vinca rosea Linn, with reference to its antitumor action. Cancer Res. 20, 1023–1031.

2 Perez, E. A. 2009 Microtubule inhibitors: Differentiating tubulin-inhibiting agents based on mechanisms of action, clinical activity, and resistance. Mol Cancer Ther. 8, 2086–2095. (10.1158/1535-7163.MCT-09-0366)

3 Jordan, M. A. 2002 Mechanism of action of antitumor drugs that interact with microtubules and tubulin. Curr Med Chem Anticancer Agents. 2, 1–17. (10.2174/1568011023354290)

4 Kundranda, M. N., Niu, J. 2015 Albumin-bound paclitaxel in solid tumors: clinical development and future directions. Drug Des Devel Ther. 9, 3767–3777. (10.2147/DDDT.S88023)

5 Jain, S., Kapoor, G. 2010 Chemotherapy in Ewing’s sarcoma. Indian J Orthop. 44, 369–377. (10.4103/0019-5413.69305)

6 Delattre, O., Zucman, J., Plougastel, B., Desmaze, C., Melot, T., Peter, M., Kovar, H., Joubert, I., de Jong, P., Rouleau, G., et al. 1992 Gene fusion with an ETS DNA-binding domain caused by chromosome translocation in human tumours. Nature. 359, 162–165. (10.1038/359162a0)

7 Jeon, I. S., Davis, J. N., Braun, B. S., Sublett, J. E., Roussel, M. F., Denny, C. T., Shapiro, D. N. 1995 A variant Ewing’s sarcoma translocation (7;22) fuses the EWS gene to the ETS gene ETV1. Oncogene. 10, 1229–1234.

8 Peter, M., Couturier, J., Pacquement, H., Michon, J., Thomas, G., Magdelenat, H., Delattre, O. 1997 A new member of the ETS family fused to EWS in Ewing tumors. Oncogene. 14, 1159–1164. (10.1038/sj.onc.1200933)

9 Delattre, O., Zucman, J., Melot, T., Garau, X. S., Zucker, J. M., Lenoir, G. M., Ambros, P. F., Sheer, D., Turc-Carel, C., Triche, T. J., et al. 1994 The Ewing family of tumors--a subgroup of small-round-cell tumors defined by specific chimeric transcripts. N Engl J Med. 331, 294–299. (10.1056/NEJM199408043310503)

10 Azuma, M., Embree, L. J., Sabaawy, H., Hickstein, D. D. 2007 Ewing sarcoma protein ewsr1 maintains mitotic integrity and proneural cell survival in the zebrafish embryo. PLoS One. 2, e979. (10.1371/journal.pone.0000979)

11 Park, H., Turkalo, T. K., Nelson, K., Folmsbee, S. S., Robb, C., Roper, B., Azuma, M. 2014 Ewing sarcoma EWS protein regulates midzone formation by recruiting Aurora B kinase to the midzone. Cell Cycle. 13, 2391–2399. (10.4161/cc.29337)

12 Park, H., Kim, H., Hassebroek, V., Azuma, Y., Slawson, C., Azuma, M. 2021 Chromosomal localization of Ewing sarcoma EWSR1/FLI1 protein promotes the induction of aneuploidy. J Biol Chem. 296, 100164. (10.1074/jbc.RA120.014328)

13 Kim, H., Park, H., Schulz, E. T., Azuma, Y., Azuma, M. 2023 EWSR1 prevents the induction of aneuploidy through direct regulation of Aurora B. Front Cell Dev Biol. 11, 987153. (10.3389/fcell.2023.987153)

14 Embree, L. J., Azuma, M., Hickstein, D. D. 2009 Ewing sarcoma fusion protein EWSR1/FLI1 interacts with EWSR1 leading to mitotic defects in zebrafish embryos and human cell lines. Cancer Res. 69, 4363–4371. (10.1158/0008-5472.CAN-08-3229)

15 Schaedel, L., John, K., Gaillard, J., Nachury, M. V., Blanchoin, L., Thery, M. 2015 Microtubules self-repair in response to mechanical stress. Nat Mater. 14, 1156–1163. (10.1038/nmat4396)

16 Rai, A., Liu, T., Katrukha, E. A., Estevez-Gallego, J., Manka, S. W., Paterson, I., Diaz, J. F., Kapitein, L. C., Moores, C. A., Akhmanova, A. 2021 Lattice defects induced by microtubule-stabilizing agents exert a long-range effect on microtubule growth by promoting catastrophes. Proc Natl Acad Sci U S A. 118, (10.1073/pnas.2112261118)

17 Igaev, M., Grubmuller, H. 2020 Microtubule instability driven by longitudinal and lateral strain propagation. PLoS Comput Biol. 16, e1008132. (10.1371/journal.pcbi.1008132)

18 Muhlethaler, T., Gioia, D., Prota, A. E., Sharpe, M. E., Cavalli, A., Steinmetz, M. O. 2021 Comprehensive Analysis of Binding Sites in Tubulin. Angew Chem Int Ed Engl. 60, 13331–13342. (10.1002/anie.202100273)

19 Andreu-Carbo, M., Egoldt, C., Velluz, M. C., Aumeier, C. 2024 Microtubule damage shapes the acetylation gradient. Nat Commun. 15, 2029. (10.1038/s41467-024-46379-5)

20 Xu, Z., Schaedel, L., Portran, D., Aguilar, A., Gaillard, J., Marinkovich, M. P., Thery, M., Nachury, M. V. 2017 Microtubules acquire resistance from mechanical breakage through intralumenal acetylation. Science. 356, 328–332. (10.1126/science.aai8764)

21 Dimitrov, A., Quesnoit, M., Moutel, S., Cantaloube, I., Pous, C., Perez, F. 2008 Detection of GTP-tubulin conformation in vivo reveals a role for GTP remnants in microtubule rescues. Science. 322, 1353–1356. (10.1126/science.1165401)

22 Aumeier, C., Schaedel, L., Gaillard, J., John, K., Blanchoin, L., Thery, M. 2016 Self-repair promotes microtubule rescue. Nat Cell Biol. 18, 1054–1064. (10.1038/ncb3406)

23 Triclin, S., Inoue, D., Gaillard, J., Htet, Z. M., DeSantis, M. E., Portran, D., Derivery, E., Aumeier, C., Schaedel, L., John, K., et al. 2021 Self-repair protects microtubules from destruction by molecular motors. Nat Mater. 20, 883–891. (10.1038/s41563-020-00905-0)

24 Manka, S. W., Moores, C. A. 2018 The role of tubulin-tubulin lattice contacts in the mechanism of microtubule dynamic instability. Nat Struct Mol Biol. 25, 607–615. (10.1038/s41594-018-0087-8)

25 Zehr, E. A., Sun, S., Sarbanes, S. L., Roll-Mecak, A. 2026 Microtubules in the axon are GDP bound but adopt a stable GTP-like expanded state. Nat Struct Mol Biol. 33, 631–640. (10.1038/s41594-026-01787-7)

26 Janke, C., Magiera, M. M. 2020 The tubulin code and its role in controlling microtubule properties and functions. Nat Rev Mol Cell Biol. 21, 307–326. (10.1038/s41580-020-0214-3)

27 Gudimchuk, N. B., McIntosh, J. R. 2021 Regulation of microtubule dynamics, mechanics and function through the growing tip. Nat Rev Mol Cell Biol. 22, 777–795. (10.1038/s41580-021-00399-x)

28 Wang, Y. L., Chen, H., Zhan, Y. Q., Yin, R. H., Li, C. Y., Ge, C. H., Yu, M., Yang, X. M. 2016 EWSR1 regulates mitosis by dynamically influencing microtubule acetylation. Cell Cycle. 15, 2202–2215. (10.1080/15384101.2016.1200774)

29 de Forges, H., Pilon, A., Pous, C., Perez, F. 2013 Imaging GTP-bound tubulin: from cellular to in vitro assembled microtubules. Methods Cell Biol. 115, 139–153. (10.1016/B978-0-12-407757-7.00010-4)

30 Hapugaswatta, H., Parrales, A., Park, H., Kim, H., Iwakuma, T., Azuma, M. 2026 The combination of EWSR1-FLI1 and loss of one EWSR1 allele leads to the induction of trisomy 8. bioRxiv. 2026.2005.2021.726567. (10.64898/2026.05.21.726567)

31 Vigano, C., von Schubert, C., Ahrne, E., Schmidt, A., Lorber, T., Bubendorf, L., De Vetter, J. R. F., Zaman, G. J. R., Storchova, Z., Nigg, E. A. 2018 Quantitative proteomic and phosphoproteomic comparison of human colon cancer DLD-1 cells differing in ploidy and chromosome stability. Mol Biol Cell. 29, 1031–1047. (10.1091/mbc.E17-10-0577)

32 Lengauer, C., Kinzler, K. W., Vogelstein, B. 1997 Genetic instability in colorectal cancers. Nature. 386, 623–627. (10.1038/386623a0)

33 Tirode, F., Laud-Duval, K., Prieur, A., Delorme, B., Charbord, P., Delattre, O. 2007 Mesenchymal stem cell features of Ewing tumors. Cancer Cell. 11, 421–429. (10.1016/j.ccr.2007.02.027)

34 Nandakumar, S., Bosche, J., Wieczorek, M., Albrecht, C. M., Konig, B., Grunewald, M., Santen, L., Diez, S., Shaebani, R., Schaedel, L. 2026 Kinesin-Induced Buckling Reveals the Limits of Microtubule Self-Repair. Adv Sci (Weinh). 13, e21721. (10.1002/advs.202521721)

35 Kalutskii, M., Grubmuller, H., Volkov, V. A., Igaev, M. 2025 Microtubule dynamics are defined by conformations and stability of clustered protofilaments. Proc Natl Acad Sci U S A. 122, e2424263122. (10.1073/pnas.2424263122)

36 Brouhard, G. J., Rice, L. M. 2018 Microtubule dynamics: an interplay of biochemistry and mechanics. Nat Rev Mol Cell Biol. 19, 451–463. (10.1038/s41580-018-0009-y)

37 Li, F., Li, Y., Ye, X., Gao, H., Shi, Z., Luo, X., Rice, L. M., Yu, H. 2020 Cryo-EM structure of VASH1-SVBP bound to microtubules. Elife. 9, (10.7554/eLife.58157)

38 Hari, M., Wang, Y., Veeraraghavan, S., Cabral, F. 2003 Mutations in alpha- and beta-tubulin that stabilize microtubules and confer resistance to colcemid and vinblastine. Mol Cancer Ther. 2, 597–605.

